# Transitioning to e-cigarettes restores the immune response to house dust mites in cigarette smoked mice

**DOI:** 10.1101/2024.12.01.626258

**Authors:** Elizabeth J. Myers, Thomas P. Huecksteadt, Noel G. Carlson, Karl A. Sanders, Kristi J. Warren

**Author notes:** Corresponding Author: Kristi J. Warren, Research Service, Building 2, GC27 500 Foothill Dr, Salt Lake City, UT 84148.

## Abstract

Since the introduction of electronic cigarettes to the US market, e-cigarettes have been posited as a safe alternative to combustible cigarettes. We developed a preclinical animal model to determine whether transitioning to e-cigarette use after up to 16 weeks of daily exposure to combustible cigarette smoke (CS) could restore normal lung immune responsiveness to house dust mites (HDM). In these studies, CS exposed animals were randomly assigned to 6 groups. (1) CS-CS mice continued combustible cigarette exposure for an additional 7 or 16 weeks, and (2) CS-recovery mice were removed from cigarette smoking where they recovered without intervention. (3) CS-carrier mice transitioned to vaporized propylene glycol (30%) with vegetable glycerol (70%) (i.e. carrier). (4) CS-salt mice transitioned to e-vapor exposure containing nicotine salt (liquid nicotine in benzoic acid + carrier), and (5) CS-base mice transitioned to daily exposures to liquid nicotine + carrier containing e-vapors. (6) Room air exposed mice, that were not smoked or exposed to e-cigarette vapors, were included as controls. We hypothesized that transitioning from CS to either of the three e-cigarette exposures (base, salt or carrier) would restore eosinophil influx into the airways following intranasal HDM administration. Here we report that shorter (7 week) e-vapor exposure containing salt, base or carrier led to significant eosinophil responses following HDM challenge. In the 16-week model, CS-base and CS-salt exposed animals did not regain their HDM responsiveness when compared to controls. CS-carrier mice did regain partial responses to HDM at 16 weeks as indicated by an increase in eosinophils compared to control mice. Lung resident lymphoid cells support the influx of eosinophils following allergen exposure. As such we measured total T cells, B cells and group 2 innate lymphoid cells (ILC2) in the lungs of each of the treatment groups. ILC2 and CD4+ T cells were reduced, and B cells were increased in the lungs of CS mice compared to controls. Numerically, the transition to nicotine-salt increased the CD3+ T cell response but transitioning to the nicotine-base significantly reduced CD19 B cells. Additional studies showed that GM-CSF protein was increased in cultured ILC and in whole lung tissues of control mice compared to CS-carrier mice indicating plasticity of the ILC2 population. RNA microarray analyses identified significant increases in GM-CSF, CCL17 and CCL24 transcripts in alveolar macrophages following the transition from CS to carrier compared to control mice. In summary, the immunosuppressive effects of CS may be restored following short-term use of e-cigarettes, but chronic use of e-cigarettes may blunt pulmonary immunity similarly to traditional cigarette smoke.

## Introduction

Electronic cigarettes, including electronic nicotine delivery systems (ENDS), were introduced to the US market in 2007, hypothetically, to aid traditional cigarette smokers with cessation.^1^ The more palatable, yet highly addictive, nicotine-salt developed by JUUL delivered higher concentrations of nicotine in e-vapors than nicotine-base in the first generation ENDS devices. The addictive nature of this nicotine-salt added significantly to the commercial success of ENDS devices leading to new, and younger, subsets of nicotine addicted individuals.^2^ In 2021, it was estimated that 4.5% of the adult population use e-cigarettes, and as most are aware, adolescent-use of vaping devices peaked in the United States in 2019 at approximately 17% of high school and middle school students combined.^3–5^ In association with a high percentage of children using these devices, as predicted by the research and medical communities, asthma rates increased in adults and adolescents in those individuals that used e-cigarettes.^6–8^

Even though the harmful effects of smoking cigarettes are well known, smoking is still the leading cause of preventable death in the United States.^8^ With 16 million people in the United States living with smoking-related disease and health complications, healthcare providers are still looking for effective ways to wean people from combustible cigarettes. In many cases individuals are not able or willing to give up nicotine completely so the switch to vaping has been proposed as a cessation technique.^9^ Generally promoting ENDS devices as a “safer” alternative to cigarette smoke for the lungs of those already diagnosed smoking related-lung disease may be accurate, but promoting e-cigarettes as safer alternative in terms of nicotine-dependence and substance abuse perhaps is not.^10^ The full impact on the lungs and overall mental health of individuals who vape will likely not be fully known for decades, especially since these products have not been around long enough for lifetime use studies. In this basic science study, our goal was to determine whether CS-induced immune suppression could be recovered in animals that transitioned from combustible cigarettes to e-cigarettes. We chose to use at least 12 weeks of CS to establish the ‘smoked lung’ effect. At that point, in contrast to never-smoked controls, no eosinophils are detected following airway allergen sensitization and challenge with HDM in CS mice thereby demonstrating significant disruption of normal immune response in the smoked lung. Following the transition to e-vapor exposures for 7 or 16 weeks, we determined whether CS-smoked animals recovered the customary allergen response to house dust mites (HDM). We found that exposure to e-cigarette vapors after CS were associated with the recovery of the normal allergen response if mice were exposed for only 7 weeks, but not if e-cigarette exposure continued to 16 weeks, suggesting ongoing inhibition of pulmonary immune responses in association with prolonged e-cigarette exposure following cigarette smoking.

## Materials & Methods

### Animal use and in vivo treatments

C57BL/6J mice were purchased from Jackson Laboratories (Bar Harbor, ME) at 8 weeks of age. After acclimation, cigarette smoked mice were exposed to a combination of mainstream and side-stream cigarette smoke from a Teague TE-10 smoking chamber (Teague Enterprises, Woodland, CA) for 225 minutes a day 5 days a week for 12-16 weeks. This dose produced 150 mg/m^3 total suspended particulates (TSP) when tested routinely. Reference cigarettes (3R4F) purchased from the University of Kentucky Agricultural Research Program were used. Untreated, age and sex-matched mice were included as controls, exposed to room air, and exposed to similar handling and noises within in the animal facility. ENDS base, salt or carrier control were delivered via Smok unit from a port on the side of the chamber. Each treatment group of mice were placed in their own dedicated subdivided Plexiglas chamber (Kent Scientific, Torrington, CT). A vacuum pump connected to the opposite side of the chamber circulated the vapor. Mice received 3 puffs (7 second puff, wait 10 seconds, repeated 3 times) twice a day. Groups of mice received carrier only, nicotine base or nicotine salt for 7-16 weeks. See below Vaping Liquid Preparation for making each vaping liquid tested in this study. These CS and ENDS exposure procedures are summarized in Figure 1A. In the last two weeks of CS or ENDS exposures, mice were sensitized with HDM (*Dermatophagoides sp*.; Greer, Lenoir, NC). (PMID: 21317378) In summary, mice were sensitized by the intranasal administration of 1 µg of HDM in 40 µL of saline (20 µl per nostril). One week later, 10 µg of intranasal HDM in saline was given (20 µl per nostril) for 5 consecutive days. Mice continued their cigarette smoke and ENDS exposures during the HDM sensitization and challenge phases of the experiment. Mice were humanely euthanized for tissue collection 96 hours after the final HDM dose. All mice were housed and used according to the *Guide for the Care and Use of Laboratory Animals* of the National Institutes of Health and in accordance with a protocol approved in advance by the Institutional Animal Care and Use Committee at the Salt Lake City VA Medical Center.

**Figure 1.**
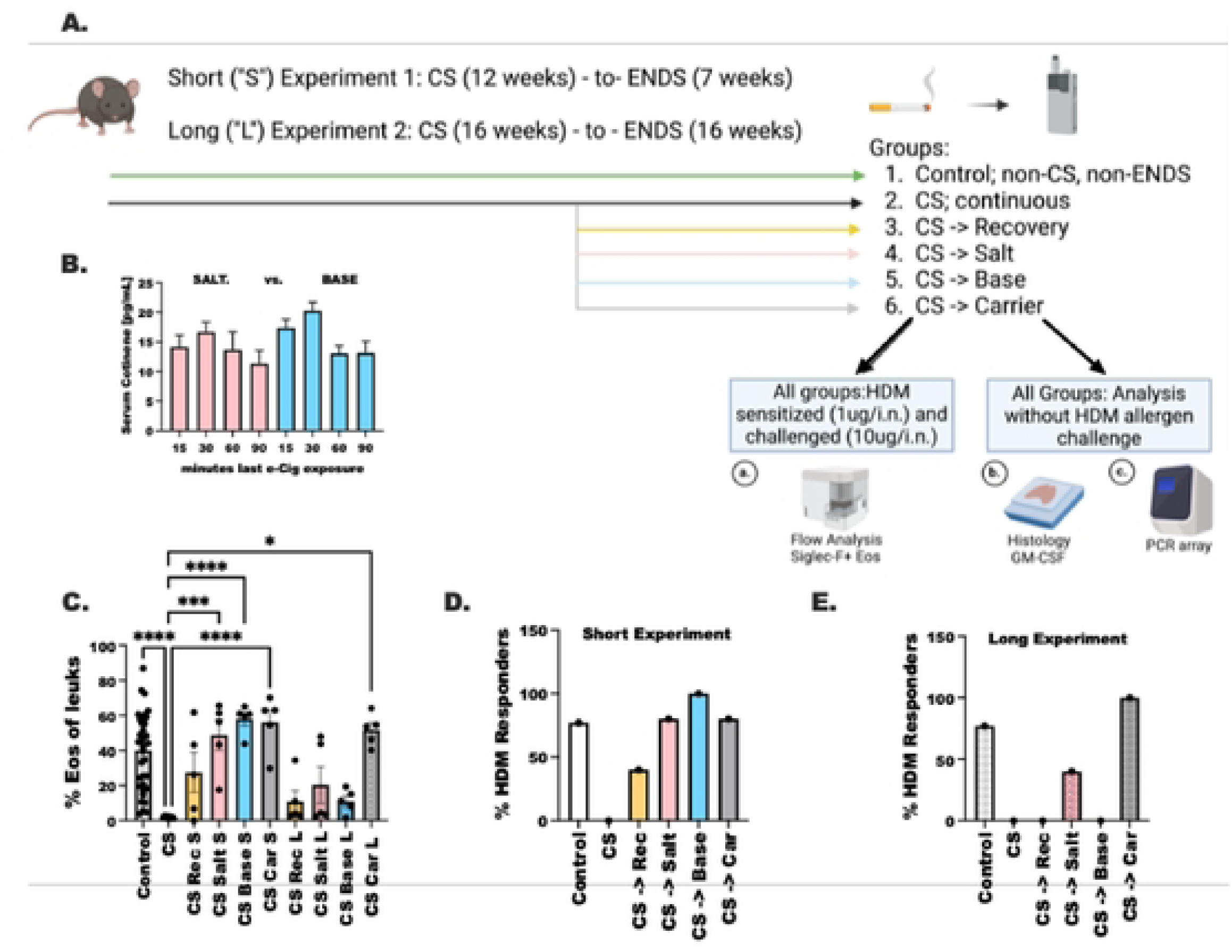
Transitioning to 7 weeks of ENDS exposures after combustible cigarette use restores a normal lung inflammatory immune response to house dust mite. Age-matched, female C57BL6J mice were exposed to fractional combustiable cigarette smoke before transitioning to ENDS. **A.** Experimental timeline for the short (12 weeks CS and 7 weeks ENDS) and long term (16 weeks CS to 16 weeks ENDS) experiments. **B.** Serum cotinine levels as measured by ELISA in mice (n-5) who were exposed to nicotine base (20 mg/mL) or nicotine salt (59 mg/mL). **C-F.** Cells present in BAL after HDM challenge were measured by How cytometry. **C.** Sigiec-F+/CD11c-cosmoph.is and **D.** Ly6C+ PMNs expressed as a percentage of the total BAL leukocyte population. Statistical analysis was performed using a one way ANOVA with a Tukey’s post-test. Statistical significance is shown as * where P <0.0001. E. Short experiment and F. long experiment expressed as percentage of mice (n -5) per treatment group with infiltrating eosinophils following HDM challenge

### Preparation of Vaping Liquids

To make the e-cigarette liquids for testing, two different solutions of nicotine (designated salt or base in this study) were combined with a carrier consisting of propylene glycol and glycerol in a 30%/70% mix which was taken from the composition of a Juul cartridge (US Patent: Nicotine salt formulations for aerosol devices and methods thereof, Juul Labs Inc.) Nicotine salt solution was made by adding benzoic acid to liquid nicotine (> 99% Sigma Aldrich St Louis MO) to a final concentration of 59 mg/ml and heated at 55° until the benzoic acid was fully dissolved. Base nicotine does not contain benzoic acid and has a final concentration of 20 mg/ml. Extensive testing was done to determine the optimal physiological concentration of nicotine delivered with either the base or salt solutions that resulted in similar plasma concentrations of cotinine, the stable metabolite of nicotine (see Figure 1B) in each group as determined by commercially available cotinine ELISA kits (Calbiotech, El Cajon, CA).

### Isolation of bronchial alveolar lavage fluid

Bronchoalveolar lavage (BAL) fluid was collected as described previously (PMID: 25803612, PMID: 23469197, PMID: 29177258). Briefly, mice received a lethal intraperitoneal tribromoethanol injection before thoracotomy and exsanguination. The trachea was exposed and a 21-guage infusion needle was inserted and secured by suture. A syringe was attached containing 1 ml of buffer (PBS, 2% FBS, 0.05% EDTA). The lung was gently flushed 5 times. The BALF was then centrifuged to pellet the cells. A fraction of the cells from each mouse were combined, incubated with CD11c coupled magnetic beads (Miltenyi biotech), and enriched with an autoMACS cell separator. The RNA was isolated and run on an RT-PCR microarray (Qiagen).

### Alveolar Macrophage (AM) cultures

Short term AM cultures were performed as described in our previous study. (PMID: 32726138) In summary, BAL was collected and pooled from groups of 5 mice and cultured for 6 hours at 37^0^ with 5% CO_2_ in the presence of mouse recombinant IL-4 (R&D systems, 404-ML) and IL-10 (R&D Systems 417-ML) at a concentration of 5 ng. PAMs AVL3288 (25 uM) (MedKoo Biosciences, Morrisville, NC), NS1738 (10uM) (Tocris, Minneapolis, MN) or PNU120596 (50 nM) (Cayman Chemical, Ann Arbor, MI) were added. Cells were collected in Qiazol (Qiagen, Germantown,MD) and flash frozen until RNA was isolated. Taqman gene expression primers were purchased from Thermo Fisher (Waltham, MA) for quantitative PCR. (PMID: 29975102 and PMID: 29117258)

### Flow cytometry

After isolation BAL cells were washed by centrifugation and resuspended in 200 uL of buffer (PBS, 2% BSA, 0.05% EDTA) on ice at a concentration of (0.5–1 × 10^6^ cells). Fc receptors were blocked and antibodies to CD45, CD11c, CD11b, Siglec-F, Ly6G, Ly6C, CD4 and/or Gr1 were added (BD biosciences, Franklin Lakes, NJ). An Accuri cell cytometer was used to collect thirty thousand live nucleated cell events for each sample. The data were analyzed using CFlow software (Ann Arbor, MI). A lymphoid panel was used to determine the numbers of T cells, B cells and group 2 innate lymphoid cells in the lung and BAL as before (Refs XX). Briefly, cells from homogenized lungs and BAL were incubated in a Zombie Aqua viability dye (eBioscience, MA) with anti-mouse antibodies against CD45, CD11b, CD11c, CD19, CD3, CD4, CD8, CD161, KLRG-1, CD127 and CD25. After fixation cellular data was acquired on the Cytek Aurora. Data were further analyzed using FlowJo v10 software.

### Lung Histology

After confirmation of euthanasia via a lethal dose of intraperitoneal tribromoethanol administration, exsanguination was performed. The diaphragm was punctured, and a small incision was made to expose the trachea into which an 18-gauge angio-catheter was inserted. The catheter was secured with suture and connected to tubing and hooked to a fluid source kept 20-25 cm above the lungs. The lungs were then rapidly filled with 10% formalin and allowed to remain filled at 20-25 cm H_2_0 pressure for 10 minutes. Lungs were carefully dissected out and placed in 10% formalin. After the lung tissue was inflation fixed the large lobe was placed in a histopathology cassette, coded and sent to the Research Histology Core Laboratory at ARUP Laboratories (Salt Lake City, Utah) for paraffin embedding, sectioning and staining.

### GM-CSF staining

To stain for GM-CSF, the Pierce Peroxidase Detection kit was used (Thermo Scientific, MA). Slides were de-parafinized according to manufacturer instructions. Citrate antigen retrieval was performed by boiling a 1x dilution of Citrate buffer pH 6.0 20X concentrate (Invitrogen, CA) and immersing the slides for 20 minutes. Slides were then washed in water and allowed to cool to room temperature. The staining protocol supplied in the kit was followed using a 1:200 dilution of the primary GM-CSF antibody (Invitrogen product number PA5-96512) and a 1:1000 dilution of the secondary antibody (Jackson ImmunoResearch product number 111-035-144). Slides were counterstained with hematoxylin. GM-CSF staining was quantitated in ImageJ using a published protocol (PMID: 31867411)

### Statistical Analysis

The long and the short experiments were analyzed using a One-way ANOVA, with a Mann-Whitney post-test to confirm between groups differences. A p-value < 0.05 was considered significant in these studies. All statistical analysis were performed on GraphPad Prism version 10.2.1.

## Results

### Transitioning to 7 weeks of ENDS exposures after combustible cigarette use restores a normal lung inflammatory immune response to house dust mites

As shown in **Figure 1A**, C57BL/6J mice were exposed to 12 or 16 weeks of combustible cigarette smoke (CS) then transitioned to electronic nicotine delivery system (ENDS) generated vapor exposures for either 7 weeks or 16 weeks respectively. For comparison, we are calling these the short (CS: 12 weeks -> ENDS: 7 weeks) and the long (CS: 16 weeks -> ENDS: 16 weeks) CS-to-ENDS experiments. The two nicotine preparations used in the ENDS groups, nicotine-salt and nicotine-base were confirmed to produce similar levels of serum cotinine despite the different concentrations of nicotine present in solution at the time of vapor generation (**Figure 1B**). The numbers of eosinophils infiltrating into the airways following HDM sensitization and challenge were measured in all treatment groups for the short and long experiments. Non-smoked and non-vaped control mice had 40% eosinophils infiltrating into the lung whereas cigarette smoke (CS) exposed animals had no eosinophil recruitment as compared to control mice (P < 0.0001) (**Figure 1C**).^11–13^ When CS mice were transitioned to non-smoking (i.e. recovery), or any of the ENDS delivered preparations (nicotine-salt, nicotine-base or carrier) for 7 weeks (in the “short” experiment), we observed significant recovery of the number of eosinophils in BAL fluid with HDM sensitization and challenge. However, in the animals that transitioned to ENDS use for 16 weeks (in the “long” experiment), the ENDS delivered nicotine-salt and nicotine base groups showed reduced eosinophil response to HDM sensitization and challenge compared to never-smoked controls. We also measured neutrophil recruitment into the airways in response to HDM. Neutrophils increased in response to HDM in the same pattern as eosinophils in the short experiment, but the responses diminished with the longer exposure CS and transition to ENDS nicotine-salt or nicotine-base in comparison to CS-alone (Supplementary Figure 1). We then compared the number of responders to HDM (mice with measurable amounts of infiltrating eosinophils) within each group in the short and long experiments (**Figure 1D and 1E**). We confirmed that in the long experiment most of the mice in the continuous CS, CS-recovery, CS-salt and CS-base groups did not recover their HDM responsiveness; only carrier induced a significant HDM response in both the short and the long e-vapor transition experiments. These results indicate that there is a window of recovery of the immune response to HDM based on the duration of CS and transition to ENDS.

### Transcriptional profile of chemokines is reduced in alveolar macrophages from mice that convert from CS to ENDS for greater than 16 weeks

To examine the effects of CS to ENDS transition in macrophage gene expression, BAL was collected from mice that transitioned from both the short and long CS to ENDS experiments. CD11c+ cells were enriched with magnetic bead sorting. RNA was isolated then applied to an RT^2^ profiler PCR array plate and data from the short (**Figure 2A**) and long **(Figure 2B**) experiments were compiled into heatmaps with clustering for the 84 genes evaluated. Different patterns of chemokine and inflammatory cytokine expression were observed in these studies. Of note, GM-CSF transcripts (Csf2, indicated with a red arrow, **Figure 2A**) were elevated in the short experiment in comparison to the all the other groups. In the long experiment GM-CSF in the CS-Salt groups was comparable to control group. Next, we determined fold-change compared to the control CD11c+ cells and found that CS-Salt exposed animals tended to express higher levels of (**Figure 2C**) CCL12, (**2D**) CCL2, (**2E**) M-CSF, (**2F**) CCL7 and (**2G**) CCL24. CXCL3 was modestly increased in the short experiment, and greatly reduced by continuous CS in the long experiment (**Figure 2H**). Notably, when we compared chemokines between the short and long experiments we find similar patterns, whereby chemokines were elevated in the short transition, but defective in the long transition in CD11c+ BAL cells (**Figure 2E-2H**). Finally, two transcripts that play a role in lymphoid cell recruitment (CCL17) and alveolar structure and health (GM-CSF) were lower in the short experiment, but contrastingly, were elevated in the long experiment. Increases were seen in CCL17 in the CS-recovery, CS-Salt, CS-Base, and to the greatest extent CCL17 increased in the CS-Carrier mice compared to continuous CS exposed mice. GM-CSF was significantly induced in the long exposure experiment, although, somewhat surprisingly, CS-Salt and CS-base had less GM-CSF transcripts as compared to CS and CS-Recovery.

**Figure 2.**
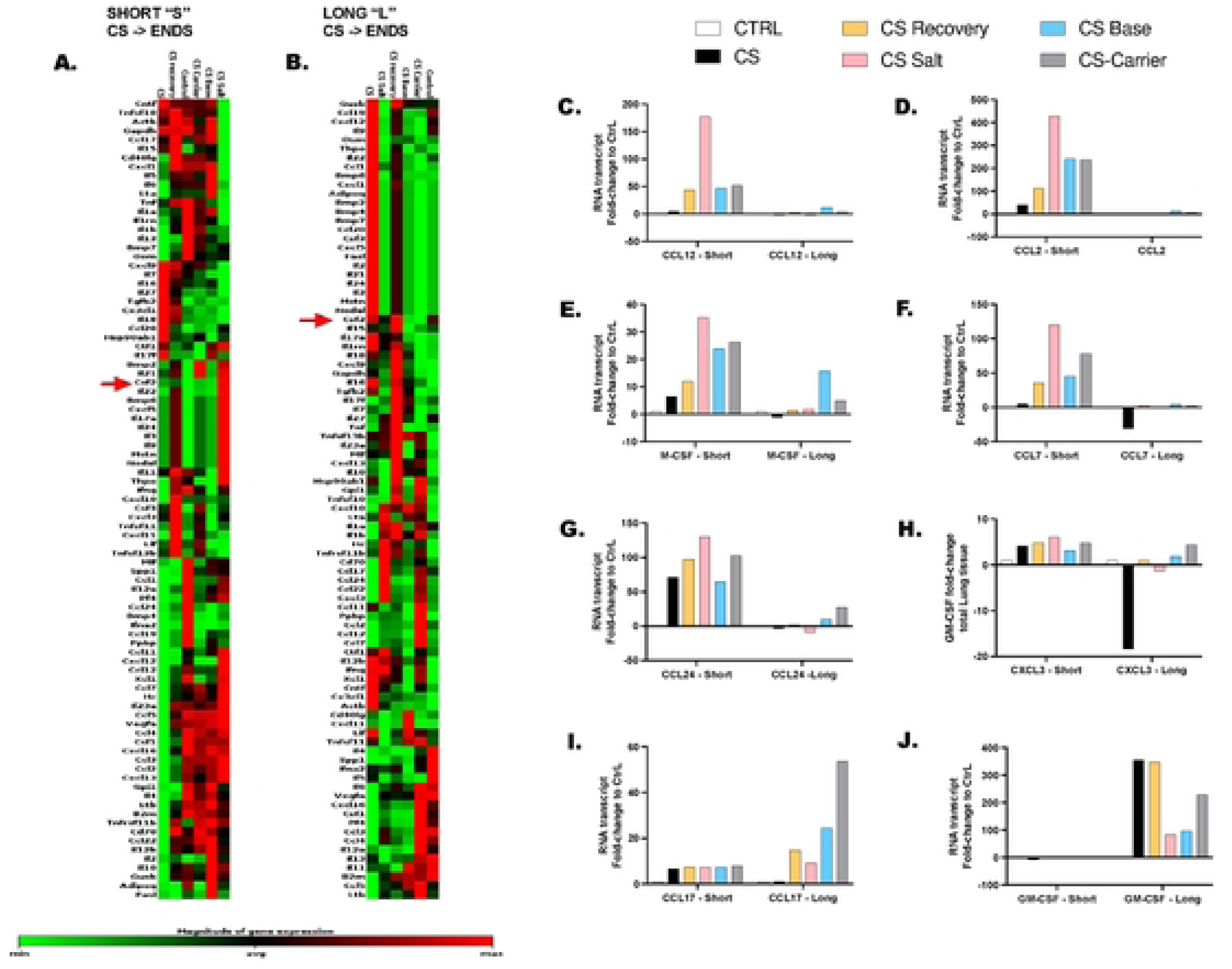
Transcriptional profile of chemokines are reduced in alveolar macrophages from mice that convert from CS to ENDS for greater than 16 weeks. Mice were treated as in Figure 1A followed by sensitization and challenge with house dust mites (HDM). BAL was collected and CD 11 c+ alveolar macrophages wore isolated for RNA PCR array analysis A. and B Dendrograms were created to compare the magnitude of transcript express across the 84 genes examined in the (A) short and the (B) long experiment. Red arrow indicates GMCSF transcripts (csf2). Quantitative PCR was used to measure transcript expression for C. CCL12. 0. CCL2. E. MCSE F. CCL7. G CCL24. H. C×CL3. I.CCL17. and J. GM CSF CD11c cells were comb nod from 5 animals/group.

### GM-CSF is suppressed across lung tissue by the conversion from CS to e-cigarette containing nicotine-salt compared to control mice

Because of the changes in GM-CSF transcript expression in alveolar macrophages in the long experiment, we used immunohistochemistry to examine expression of this protein across the lungs from mice in each treatment group. Lung tissues were scored for general inflammation around the vasculature, airways and alveoli for each of the treatment groups from the long experiment (**Figure 3A, 3B**). In contrast to the results presented in Figure 1, the animals in these experiments were not sensitized or challenged with HDM before analysis, and correspondingly exhibited only a moderate increase in inflammation noted with CS as compared to the control group. In this evaluation we found that CS-recovery and CS-salt had significantly less inflammation when compared to CS treatment alone. That is to say that the inflammation was decreasing, and more comparable to the non-smoked, non-ENDS exposed control animals, than with the CS exposed animals. GM-CSF is a pleiotropic cytokine that is associated with lung and alveolar health. We assessed GM-CSF in an unbiased manner to determine whether total expression of this protein was modulated in the various treatment groups (**Figure 3C, 3D**). We show that GM-CSF was reduced in the CS-Salt exposed animals when compared to all other groups.

**Figure 3.**
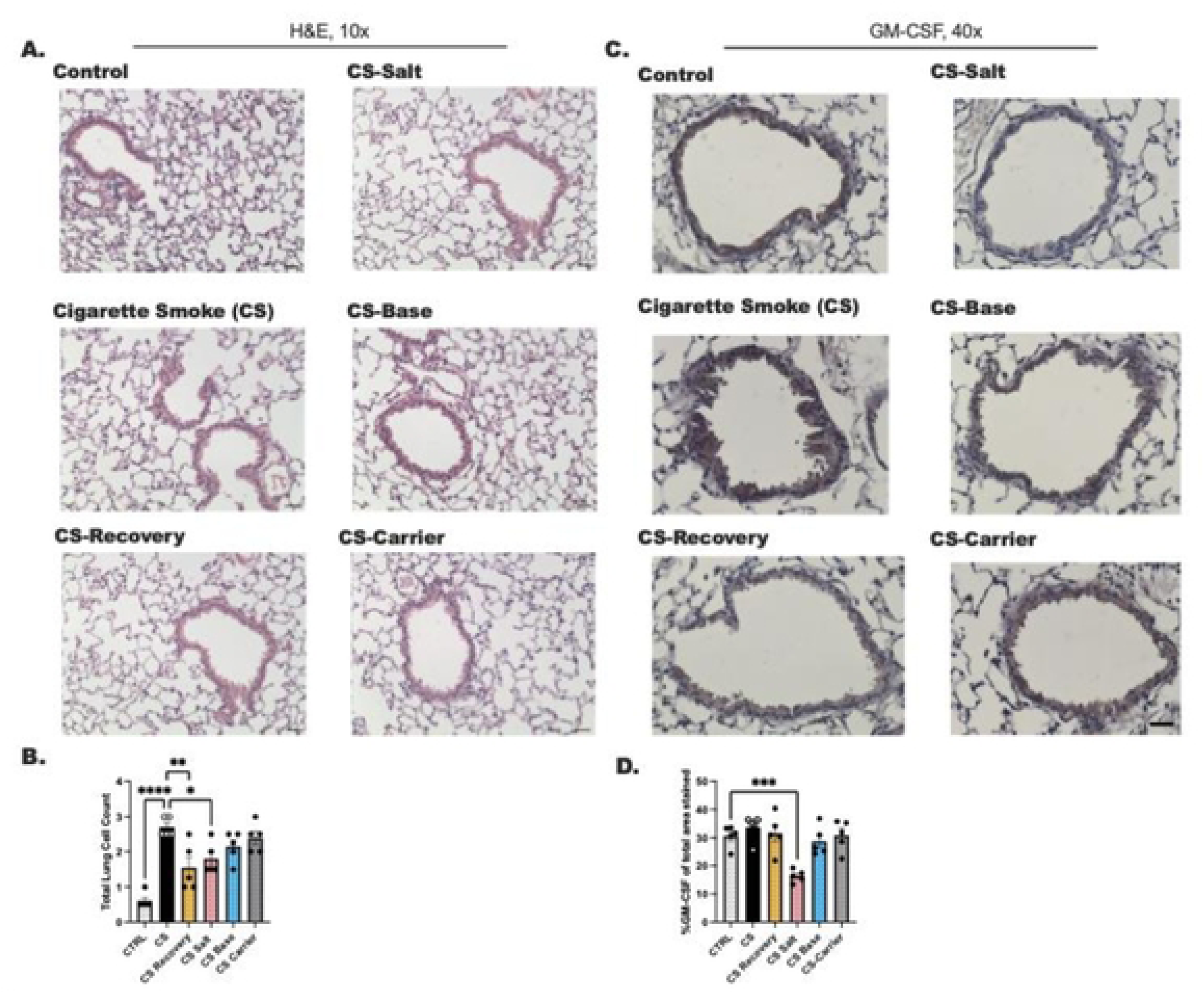
GM-CSF is suppressed across lung tissue by the conversion from CS to e-cigarette containing nicotine-salt compared to control mice. Animals were prepared as shown in Figure 1A according to the long exposure protocol. Lung tissue was formalin fixe**d** under constant pressure in the chest cavity then excised. Lungs were paraffin embedded and sectioned at 8 microns followed by staining with H&E (**2A,2B**) ant-GM-CSF with peroxidase labeling and eosin counterstain (**2C,2D**). Inflammation was scored by a pathologist that was blinded to the treatment conditions. Airway and vascular inflammation is reported as one value in 2B Fiji image J was used to calculate the percent of GM-CSF staining per total area (n=5); 20. At least 2 sections were evaluated per mouse, with a total of 5 animals evaluated per group across two experiments. Statistical analysis was performed using a one-way ANOVA with a Kruskal-Wailis post-test Statistical significance is shown as *** where P < 0.001. ** P < 0.01 and * P < 0 05. Scale bar represents 50 uM in images.

### Type 1 and type 2 positive allosteric modulators of the nicotinic acetylcholine a7 receptors suppress CCL24 transcript expression in alveolar macrophages

We have presented data showing the effects of CS and ENDS on expression of several genes. However, an important question is how nicotinic receptors may contribute to these changes. We approached this question using pharmacological manipulation of the nicotinic *⍺*7 receptors in alveolar macrophages to focus on whether nicotine actions on *⍺*7 contributes to the changes in GM-CSF (Figure 3) and chemokines (Figure 2) that modulate eosinophil recruitment following allergen challenge. Nicotine can act initially as an agonist of *⍺*7, but then rapidly desensitizes the *⍺*7 receptor upon high exposures to nicotine. Thus, in e-cigarette exposures high-levels of vaporized-nicotine are an antagonist of *⍺*7. In our previous studies, positive allosteric modulators (PAMs) of the *⍺*7 nicotinic acetylcholine receptor counteract this desensitizing effect. PAMs can reactivate *⍺*7 by increasing agonist-induced currents and inhibit the desensitization of the *⍺*7 receptor.^14^ We have found that PAMs can restore eosinophil recruitment following HDM challenge that had been suppressed by aerosolized nicotine.^13^ Upon evaluating our earlier studies, we wanted to determine whether nicotine actions through the *⍺*7 nicotinic acetylcholine receptor ultimately desensitizes cholinergic signaling, leading to suppressed chemokine production in alveolar macrophages, which could then be restored through the addition of one of the PAMs pharmacologic therapies. In these next experiments animals were exposed acutely (1-3 weeks) or chronically (7-8 weeks) to ENDS generated vapors containing nicotine-salt, nicotine-base or carrier alone. BAL cells were collected from those animals then cultured for 6 hours as before with IL-4 and IL-10 (previously shown to induce CCL24, IL-1B, CCL17), and essentially serve as an *ex vivo* proxy for HDM stimulation. After 1-3 weeks of *in vivo* treatment with vaporized nicotine-salt, -base or carrier alone, IL-4/IL-10 induced a significant *in vitro* increase in CCL24 transcripts, which was then suppressed with the addition of the type 1 PAMs of NS-1738 or AVL-2388 (**Figure 4A**). This pattern of suppression was similar between all groups indicating that *in vivo* treatment with salt, base or carrier did not affect the alveolar macrophage response to *ex vivo* cytokine stimulation. The type 2 PAM, PNU-120596 had no suppressive effect on CCL24 expression in these experiments.

**Figure 4.**
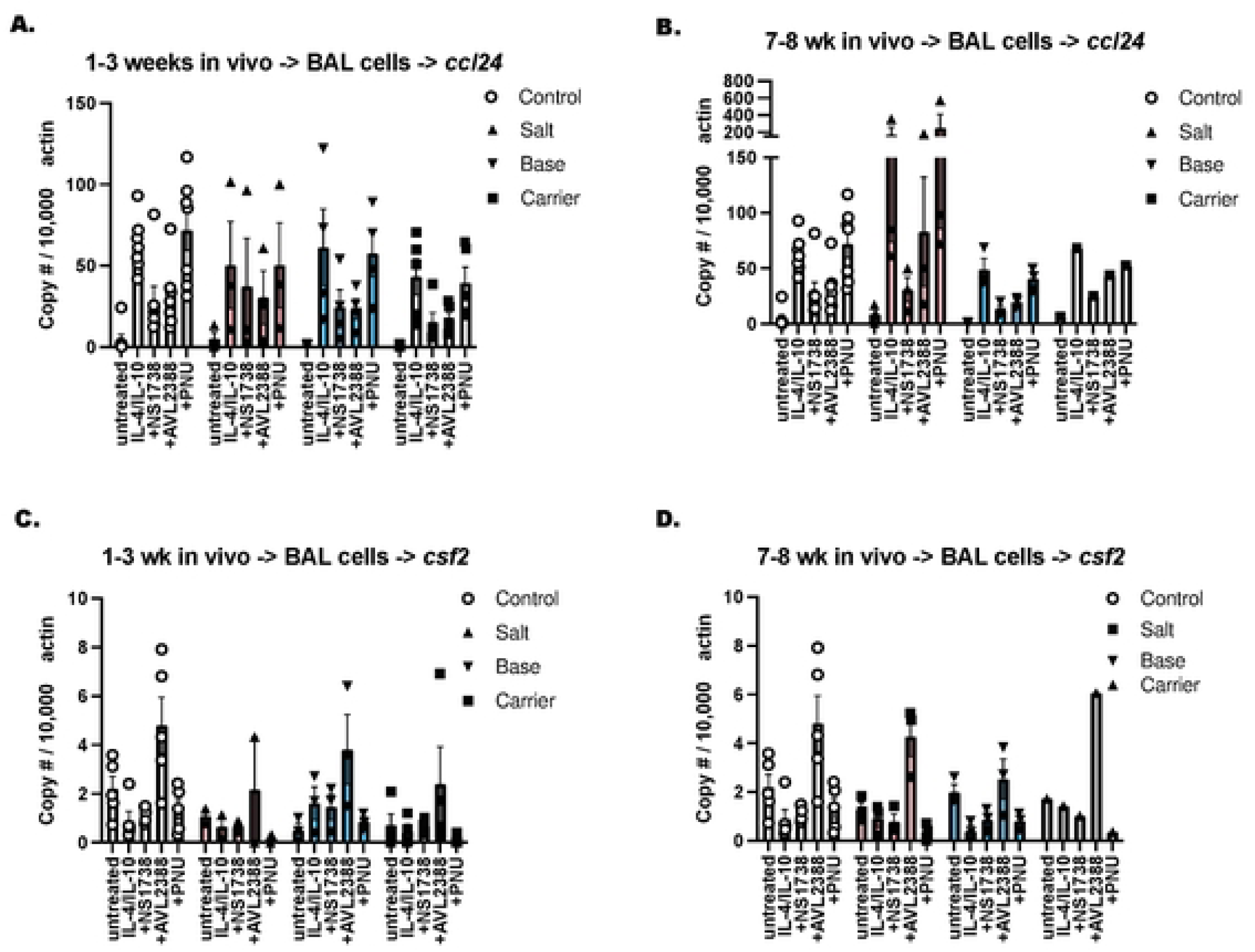
Type 1 and type 2 positive allosteric modulators of the nicotinic acetylcholine receptors suppress CCL24 transcript expression in alveolar macrophages. BAL was collected from groups of mice (n= up Io 20 mice summarized from 3-4 independent experiments) that were vaped with carrier, nicotine salt or nicoline base as well as non vaped control mice tor either 1-3 or 7-8 weeks and cultured in the presence of IL-4/10 (5ng/mL) +/-NS 1738 (10 uM).AVL3288 (25 uM) and PNU120596 (50 nM) as previously described. RNA was isolated and RT-PCR was run using Taqman primers **to** CCL24 (**A and B**) or GM CSF (CSf2; **C and D**) Graphs (**A-D**) show CCL24 or CSF2 gene expression as compared to copy number of B actin calculated using the delta delta Ct method.

At 7-8 weeks of *in vivo* exposure to salt (**pink bars, Figure 4B**) there was a significant increase in IL-4/-10 induced CCL24 transcript expression that was similarly suppressed by NS-1728 and AVL-2388, but again, no effect was noted for the type 2 PAM of PNU 120596 in the IL-4 and IL-10 responsiveness of the BAL macrophages in these experiments. Although alveolar macrophages are only a modest source of GM-CSF we were surprised to note that AVL-2388 promoted transcript expression of GM-CSF in all acute (**Figure 4C**) and chronic (**Figure 4D**) *in vivo* treatment groups when combined with *ex vivo* IL-4/IL-10 cytokine stimulation. Collectively, our results indicate that type 1 PAMs, particularly AVL2388, may be an effective strategy for restoring GM-CSF in the alveolar spaces as that loss of GM-CSF in CS-Salt exposed animals may promote problems that would be associated with a GM-CSF deficiency. Those problems include the accumulation of mucins, improper surfactant protein production and clearance, and poorer gas exchange for animals that converted from CS to nicotine salt.

### Lymphoid cell populations are differentially modified by the transition from CS-to-ENDS

We next assessed whether there were notable changes in the different lymphoid cell types in CS-to-ENDS mice challenged with HDM in the long exposure model. Lung and BAL CD45+ cells were evaluated in each of the treatment groups of mice prepared as before with HDM sensitization and challenge. We first determined whether the proportions of live versus dead CD45+ cells were different between groups (**Supplemental Figure 2**). We found that Control mice had the greatest number of live CD45+ cells compared to CS exposed animals. CS-base and CS-carrier mice trended towards having significantly more live CD45+ cells as compared to those mice that continued cigarette smoking. There were no differences in the numbers of dead cells detected when comparing the groups when lymphoid cells were examined. Not surprisingly, CD3+ T cells (**Figure 5A**), CD4+ T cells (**Figure 5C**), and ILC2 (**Figure 5D**) were suppressed in the CS mice compared to control mice. B cells, however, were increased in CS mice compared to the non-smoked, non-vaped control mice (**Figure 5B, 5F**). Total CD3+ T cells were increased in the CS-Salt mice compared to CS mice when the percentage of CD45+cells (**Figure 5A**) were determined, and when the absolute counts (**Figure 5E**) of this population were compared between these two groups. The percentage of B cells in the CS mice compared to CS-Base exposed animals were significantly reduced (**Figure 5B and 5F**). CD4+ T cells were increased in the CS-Carrier mice compared to CS mice. Lung ILC2 numbers were reduced by cigarette smoking (**Figure 5D, 5H**) in comparison to untreated controls. The transition to recovery, nicotine-salt, nicotine-base or carrier alone did not increase the numbers of ILC2 significantly above the levels measured in the CS mice.

**Figure 5.**
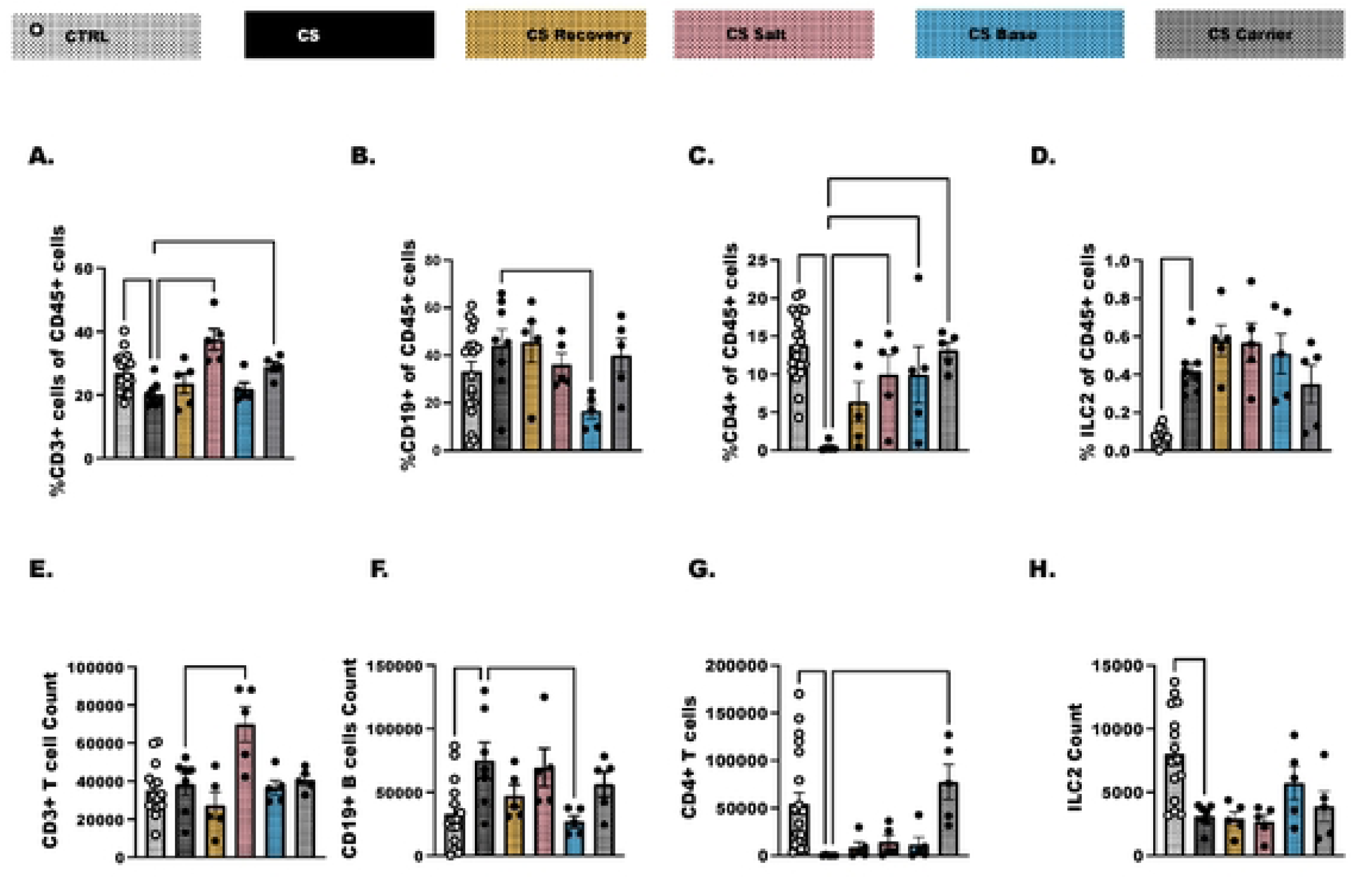
An assessment of lymphoid populations following the transition from CS to ENDS after allergen challenge. Lung tissue was collected from a subset of mice to assess the lymphoid cells that had infiltrated into the airways and lungs following HOM challenge after mice had transitioned from CS to ENDS-salt. * base or -carrier. CS only mice and animals that quit smoking (recovery) were included as controls. The frequencies and absolute counts of total (A, E) CD3+ T cells. (B, F) CD19+ B cells, (C. G) CD3+CD4+ T cells, and (D, H) group 2 innate lymphoid cells were determined by flow cytometry. These data are representative of 4 separate experiments where control mice (n=20) and CS (n=10) mice were included each day as controls. Sample size is n=5 for other treatment groups. One-way ANOVA was used to determine differences between groups, with Dunnett’s multiple comparison test to determine whether treatment groups were different that untreated controls.

### Innate lymphoid cells are responsive to activation and release different cytokine based on which treatment group from which they are derived

We next isolated innate lymphoid cells from the lungs of these animals and stimulated with a cytokine mixture that supports innate lymphoid cell survival (IL-2 and IL-7) and cytokine production (IL-33, airway-derived cytokine). We and others have shown activation of ILC2 with these cytokines previously.^15–17^ The goal was to determine whether ILC responsiveness was restored in any of the mice that transitioned away from cigarette smoking to ENDS delivered nicotine-salt, -base, or carrier (**Figure 6A-6G**). The ILC activation experiments yielded a unique profile depending on which treatment group the cells were isolated from. IL-17 production was decreased in the control mice, CS-recovery mice, and CS-salt exposed mice as compared to the CS mice (**Figure 6A**). CCL22, GM-CSF and CCL3 were significantly elevated in ILC that were enriched and activated in the CS-recovery mice compared to the continuous CS mice (**Figure 6B-D**). Next, we detected a significant production of CCL5, CCL2, and CCL11 from ILC isolated from the lungs of CS-Carrier mice compared to those ILC isolated from the CS mice (**Figure 6E-G**). Taken together this indicates that lymphoid cells are differentially modified by the components of e-liquids in ENDS devices. Lung derived ILC specifically are an immune population that are highly responsive to airway triggers; they may represent important cellular targets of therapies for individuals that experience an airway injury associated with e-cigarette use.

**Figure 6.**
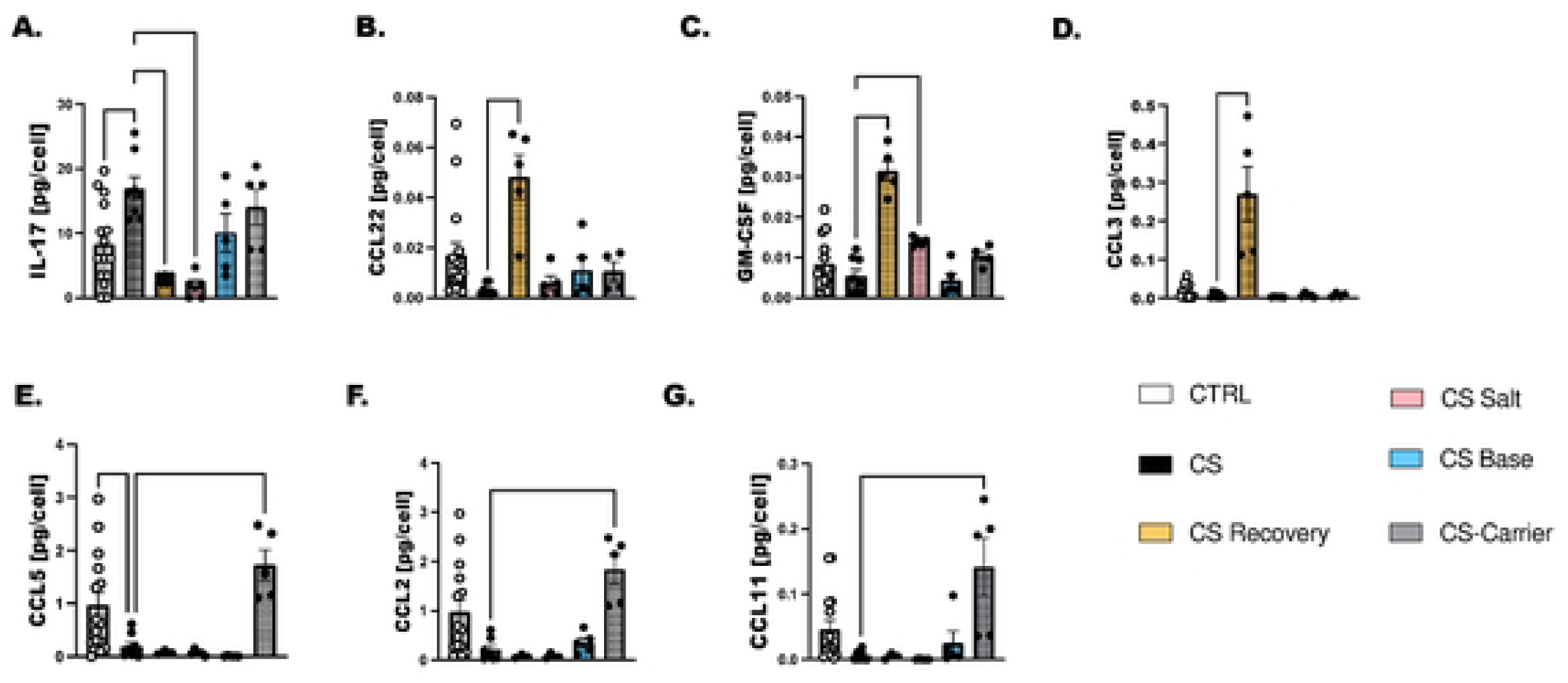
Total innate lymphoid cells (LC) were enriched from the lungs and cultured for 4 days with a cytokine cocktail of survival cytokines (IL-2. IL-7; both at 10 ng/mL) and the airway derived cytokine (IL-33; 10 ng/mL). Cytokine levels were determined using a customized legend plex assay to analyze 25 ul of cell culture media. A. IL-17. (B) CCL22. (C) GM-CSF. (D) CCL3. (E) CCL5. (F) CCL2. and (G) CCL11. As before, these data are representative of 4 separate experiments where control mice (n=20) and CS (n=1O) mice were included each day as controls. Sample size is n=5 per treatment group. One-way ANOVA was used to determine differences between groups, with Dunnett’s multiple comparison test to determine whether treatment groups were different that untreated controls.

## Discussion

These studies were initiated to evaluate whether the transition from cigarette smoking (CS) to electronic nicotine delivery systems (CS-to-ENDS) is beneficial to lung health in comparison to just continuing smoking. In previous publications,^12^ 12 weeks of cigarette smoke (CS) significantly reduced eosinophil recruitment to the airways following challenge with HDM. Others have similarly shown that CS suppresses immunity, which explains the tendency for cigarette smoking individuals to have a deficient immune response to respiratory bacterial and viral infection.^18, 19^ HDM sensitization and challenge represented a stable, well-characterized immune response that we could repetitively perform to answer significant questions for the research community regarding whether e-cigarettes are better for pulmonary health as compared to continuous smoking. To our knowledge this is the first experiment of its kind that characterizes the basic science behind CS-to-ENDS exposure over both short and longer periods of time. These models represent someone transitioning to e-cigarettes to quit smoking (‘short’ model), or a longer period of exposure that represents to us someone that transitioned to e-cigarettes without quitting (‘long’ model).^20^ In the UK, it is public policy to encourage smokers to switch to e-cigarettes even if individuals can’t achieve the ‘A’ standard of smoking cessation.^21^ This has been a successful strategy for moving high-risk individuals, i.e. those with COPD, asthma, cardiopulmonary complications, away from detrimental combustible cigarette use. Cigarette smoking is still the leading cause of preventable deaths in the US and UK. We show here that there are certainly unique immune characteristics related to the whole lung landscape and alveolar macrophages that arise from the CS-to-ENDS transition.

In earlier studies, before vaping devices were invented, adult mice were exposed to aerosolized nicotine base in water for up to 12 weeks. Nicotine significantly reduced eosinophil recruitment following HDM exposure,^13^ that was not dissimilar from the study showing suppression with CS. In the nicotine studies, hyperactivation of the *⍺*7 nicotinic acetylcholine receptor with nicotine essentially desensitized cholinergic signaling tied to a normal immune response. An important distinction with ENDS is that e-vapors contain higher concentration and a purer form of nicotine compared to burned cigarette smoke,^22^ these vapors penetrate deeply into the lung reaching the alveolar spaces more efficiently than combustible cigarette smoke.^23^ In the current study, we compared two formulations of nicotine that we confirmed were similarly metabolized in animals. Nicotine-salt is the most widely used form of nicotine in e-liquids vaporized in ENDS devices.^24^ The nicotine salt was prepared and tested following the patent from JUUL which is proving more tolerable than nicotine base. In our hands, nicotine-base, at a concentration above 20 mg/mL was toxic to animals. Comparatively, 20 mg/mL of nicotine-salt is the maximum amount of nicotine that can be legally sold in UK. This level of nicotine is suggested for those heavy smokers that are attempting to transition to e-cigarettes. Our nicotine-salt was constituted with the humectants, propylene glycol and vegetable glycerin, and we included a PG/VG only control in these studies. Our data show that each component altered the response to HDM challenge, with the most pro-allergy response mounted predominantly with the humectants alone. We are not the first to show an enhanced immune or inflammatory response with PG/VG,^25^ however, we anticipated that prior exposure to CS would blunt part of that PG/VG hypersensitivity response to HDM. This was not the case as the transition from CS to Carrier in both the short and long experiments yielded a significant eosinophil response. The carrier data highlights that the lungs can still be restored to normalcy if the proper stimulus is applied, as long as that stimulus does not contain nicotine perhaps. Innate lymphoid cells and CD4+ type 2 T helper cells are known to support a pro-allergy response, eosinophil recruitment and survival. When we evaluated lungs from the long exposure experiment, we found significantly more CD4+ T cells that unfortunately we were limited in our capacity to characterize in this study. We did however characterize the innate lymphoid cells with an *ex vivo* cell activation assay, and while group 2 innate lymphoid cell numbers were suppressed in all groups that were cigarette smoked for 16 week regardless of what they transitioned to, e.g. CS-salt, -base, or -carrier, when these cells were activated *ex vivo*, they produced CCL5 (pan-leukocyte chemoattractant), CCL2 (monocyte chemoattractant), and CCL11 (eosinophil chemoattractant) when they were isolated specifically from the CS-carrier mice. One additional striking finding from ILC culture was the production of IL-17 in the continuous CS mice, the CS-base and CS-carrier mice compared to the control treated mice. To our knowledge ILC populations that line the airway epithelial surfaces of the lungs had not been assessed in previous e-cigarette studies, and certainly not in the CS-to-ENDS model. Our report here shows ILC that were isolated from the CS-Recovery mice had ample production of CCL22 (as ILC2 chemoattractant), GM-CSF, and CCL3. The ramifications of these findings on pulmonary immunity are still under investigation, but our findings do indicate that transitioning to non-smoking and non-vaping is obviously the safest outcome for the restoration of normal immunity, but a successful restoration of immunity may have a limited window of opportunity based on duration of shorter ENDS use after transitioning away from CS.

While the limitations of the study are quite broad given the length of time it takes to prepare animals for these studies, a few are noted here. Vaping products are constantly changing, or evolving, for these lengthy studies we picked the components that were more widely used to define whether CS-to-ENDS was truly a safe alternative to continuous smoking. In 2019, when we started our studies, there were approximately 16,300 commercially available, vaping-products containing combinations of flavorings and chemicals that are still not completely characterized.^26^ We also did not experiment with voltages that would heat at different rates and vaporize e-liquids differently, and we didn’t explore the various synthetic nicotine products that are escaping FDA regulation because of slight changes to the chemical make-up. Additionally, with each new generation of ENDS systems developed (we are currently on the fourth generation), the concentration of flavorings, nicotine and humectants delivered to the lung will require additional characterization.

This highlights the challenge that the medical community faces for treating those patients that that either transition to vaping from cigarette smoking, co-use cigarettes and e-cigarettes, or those that successfully transition to e-cigarettes only. We started to work with the positive allosteric modulators of the nicotinic acetylcholine receptors because there may be potential there to restore GM-CSF production in cases where the lack of GM-CSF may be reducing alveolar health and function. Since these PAMs are under clinical trials in neurologic diseases, like schizophrenia, and Alzheimer’s, it would be potentially easier to transition these drugs to pulmonary conditions. Hypothetically, the blunted responses in the alveolar macrophages may indicate that the nicotine acetylcholine receptors might be a good target for reducing allergic responses like rhinosinusitis, or controlling asthma symptoms for those with the heightened eosinophilia, or high type 2 asthma. To our knowledge only one study evaluated the *⍺*7 nicotinic acetylcholine receptor agonist (GTS-21) in IL-33 induced allergic inflammation and showed a profound effect on reducing ILC2 activation.^27^ The evaluation of PAMs in allergic inflammation only is well outside the scope of the current study, but worth exploring potentially in future studies.

In summary, we noted that e-cigarettes could be promoted as a short-term cessation tool to help individuals quit combustible cigarettes. But, long-term use of e-cigarettes is compounded by many variables that make e-cigarette a risky alternative to CS in the long run. As we showed here, longer term use of vaping following cigarette smoke does not restore a normal lung response to an allergen but continues to suppress normal immunity.

## Acknowledgements

The authors would like to acknowledge the ARUP Research Histology Core Laboratory and the University of Utah Flow Core for their help with this project. The authors acknowledge the critical evaluation and suggestion of the manuscript by Robert Paine III.

## Author Contributions

EJM and KJW conceived and designed research; EJM, KJW, NGC and TPH performed experiments; EJM, KJW, TPH analyzed data; EJM, KJW, NGC interpreted results of experiments; KJW and EJM prepared figures; KJW and EJM drafted manuscript; EJM, KJW, and NGC edited and revised manuscript; EJM, KJW, TPH and NGC approved final version of manuscript.

---

**Supplemental Figure 1.**
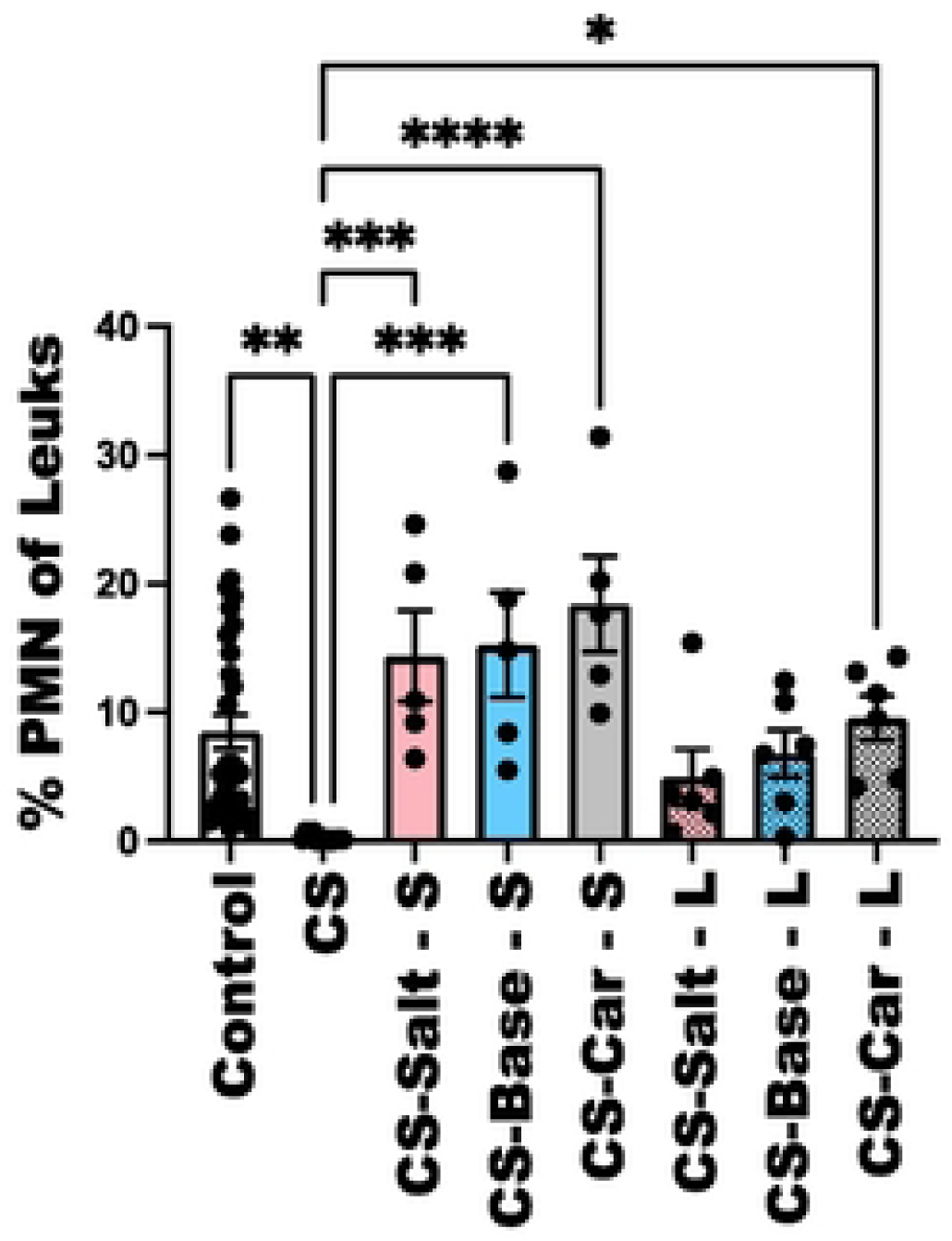
Percent PMNs in the leukocyte cell population of the BAL in the Short (S) and long (L) experiments as measure by Bow cytometry (CD4-Ly6C+CD11c+). As shown here cigarette smoked mice have a significant lack of PMNs present in the BAL which is consistent with ne fact that cigarette smoked mice do not haw infiltrating cells present in the lungs following an allergen challenge. Statistical analyses was performed using a one-way ANOVA as before (FIGURE 1) post test. Statistical significance is shown as p < 0.05.

**Supplemental Figure 2.**
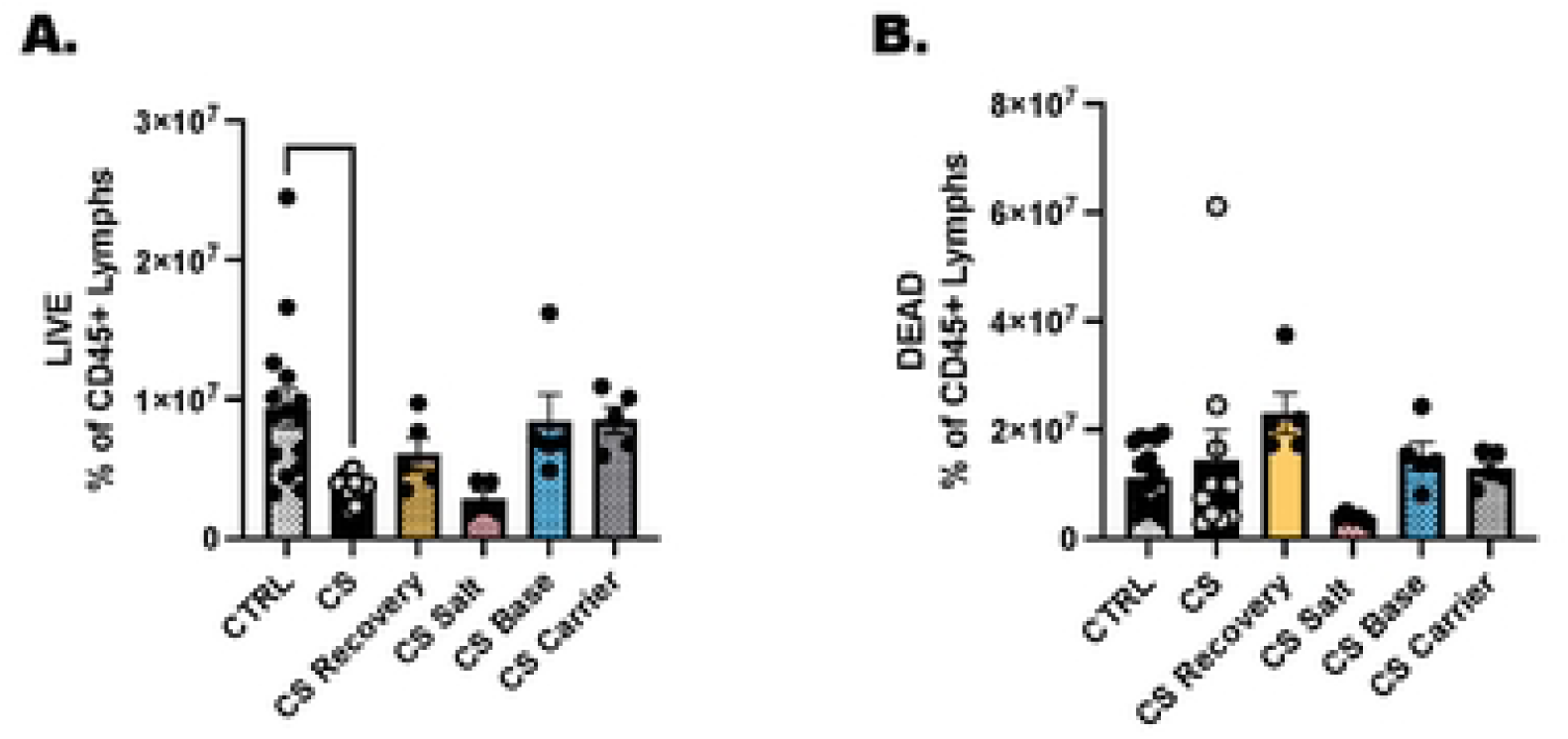
Live and dead cells were measured by flow cytometry following the various treatments in the long experiment. A Live cells were designated as CD45+ viability dye negative. B. Dead cells were designated as CD45+ and viability dye positive.

## Notes

### Competing Interest Statement

The authors have declared no competing interest.

